# Impact of *Nosema ceranae* invasion on sucrose solution consumption, midgut epithelial cell structure, and lifespan of *Apis cerana cerana* workers

**DOI:** 10.1101/2022.01.09.475580

**Authors:** Qi Long, Minghui Sun, Xiaoxue Fan, Wende Zhang, Dingding Zhou, Ying Hu, Zixin Wang, Kaiyao Zhang, Kejun Yu, Haodong Zhao, Yuemei Song, Zhongmin Fu, Dafu Chen, Rui Guo

## Abstract

*Nosema ceranae* is an intracellular fungal parasite for honeybees, leading to chronic disease named bee nosemosis with worldwide distribution. Asian honeybee (*Apis cerana*) is the original host for *N. ceranae*, but the impact of *N. ceranae* infection on *A. cerana* physiology is largely unknown. In this current work, workers of *Apis cerana cerana*, a subspecies of Asian honeybee, were artificially inoculated with *N. ceranae* spores and reared under lab conditions, followed by detection of fungal spore load as well as host sucrose solution consumption, midgut epithelial cell structure, and lifespan. The result of spore counting suggested that the spore load in the host midgut decreased significantly during 1 dpi-2 dpi, whereas that displayed an elevated trend among 2 dpi-13 dpi. The sucrose solution consumption of workers in *N. ceranae*-inoculated groups among 1 dpi-20 dpi was always higher than that of workers in un-inoculated groups; additionally, the difference of sucrose solution consumption between these two groups at 4 dpi, 5 dpi, and 13 dpi was of significance. Based on microscopic observation of paraffin sections, darkly stained parasites were clearly detected in the midgut epithelial cells of *N. ceranae*-inoculated workers at 7 dpi-10 dpi, whereas no parasite was observed in those of un-inoculated workers. In addition, the boundaries of un-inoculated host epithelial cells were intact and the darkly stained nucleus were clear, while the boundaries of midgut epithelial cells of *N. ceranae*-inoculated workers were blurred, the nucleus were almost disappeared, and the nucleic acid substances were diffused. Moreover, the survival rates of workers in both *N. ceranae*-inoculated groups and un-inoculated groups at 1 dpi-5 dpi were pretty high and then started to decrease at 5 dpi; the survival rate of workers in *N. ceranae*-inoculated groups was always lower than that in un-inoculated groups, with significant difference between these two groups during 11 dpi-20 dpi. These results together indicate that the quantity of fungal spores continuously elevated with the microsporidian multiplication, causing energetic stress for workers and host cell structure damage, which further negatively affected the host lifespan. Our findings offer a solid basis not only for exploring the molecular mechanism underlying *N. ceranae* infection but also for investigating the interaction between *N. ceranae* and eastern honeybee.

## 1. Introduction

*Apis cerana* is the original host for *Nosema ceranae*, a fungal parasite that results in bee nosemosis, which is a chronic disease frequently occurred in colonies throughout the world [1]. After long-term interaction and coevolution, *A. cerana* and *N. ceranae* have already adapted to each other [2]. Previous studies suggested that *N. ceranae* infection could lead to damage of mid-gut epithelial cell structure, increased consumption of sucrose solution, inhibition of apoptosis, immunosuppression, earlier foraging activity, as well as shortened lifespan of *A. mellifera* workers [3-4]. Comparatively, there are few studies associated with the influence of *N. ceranae* infestation on *A. cerana*, hindering a deeper understanding of microsporidian-eastern honeybee interaction [5-8]. Recently, our group parsed the immune response of *A. c. cerana* workers to *N. ceranae* invasion based on transcriptomic investigation, and revealed that different cellular and humoral immune responses were utilized by *A. c. cerana* and *A. m. ligustica* workers to defense against the same fungal infection [9]. In this present study, for the sake of evaluating the impact of *N. ceranae* infection on *A. c. cerana* workers, clean spores of *N. ceranae* was prepared to artificially inoculate newly emerged workers of *A. c. cerana*, followed by detecting the fungal spore load as well as host sucrose solution consumption and cumulative survival rate; additionally, paraffin sections of midgut tissues were prepared and subjected to microscopic observation. Our data will not only offer valuable experimental evidence for *N. ceranae* infection of *A. c. cerana* workers, but also contribute to further investigation of host response, parasite infestation, and host-parasite interaction.

## 1. Results

### 2.1 Inoculation of *A. c. cerana* workers with *N. ceranae* spores

Microscopic observation showed the purified spores were oval and highly refractive (**Figure 1A**). Further RT-PCR identification indicated that the signal fragment with expected size (approximately 76 bp) could be amplified from the spores, while no fragment can be amplified when using specific primers for *N. apis* (**Figure 1B**). The inoculation of single worker, artificial rearing under lab condition, and preparation of midgut sample were presented in **Figure 1C-E**.

**Figure 1.**
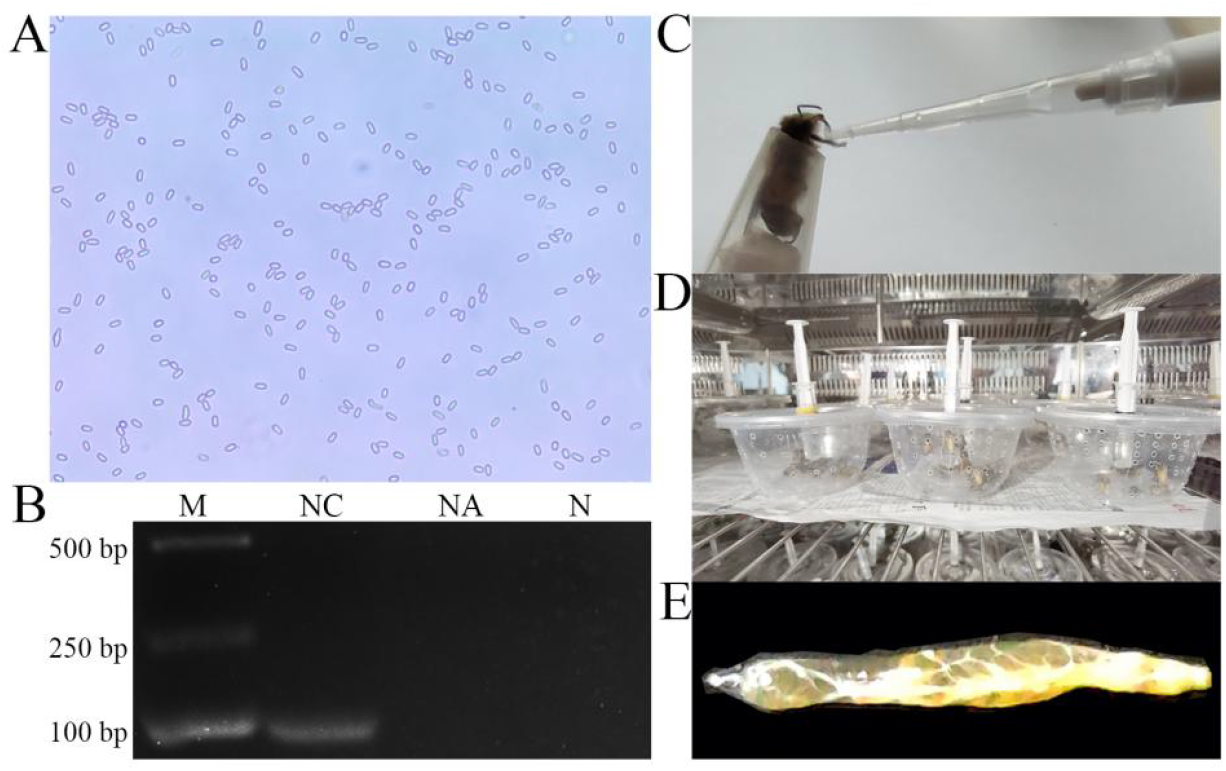
Artificial inoculation and rearing of *A. c. cerana* workers with purified spores of *N. ceranae*. (A) Microscopic observation of *N. ceranae* spores derived from Percoll discontinuous density centrifugation (400 times amplification). (B) Agarose gel electrophoresis for PCR amplification products from purified spores. Lane M: DNA marker; Lane NC: specific primers for *N. ceranae*; Lane NA: specific primers for *N. apis*; Lane N: sterile water (negative control). (C) Artificial inoculation of a fixed worker using a pipette. (D) Artificial rearing of workers kept in plastic cages in incubator. (E) A midgut tissue of a worker after dissection.

### 2.2 Dynamics of *N. ceranae* spore load in the midguts of *A. c. ceranae* workers

The result of spore counting suggested that the spore load in the host midgut decreased significantly during 1 dpi-2 dpi, whereas that displayed an elevated trend among 2 dpi-13 dpi; however, the spore load began to decline at 13 dpi **(Figure 2)**.

**Figure 2.**
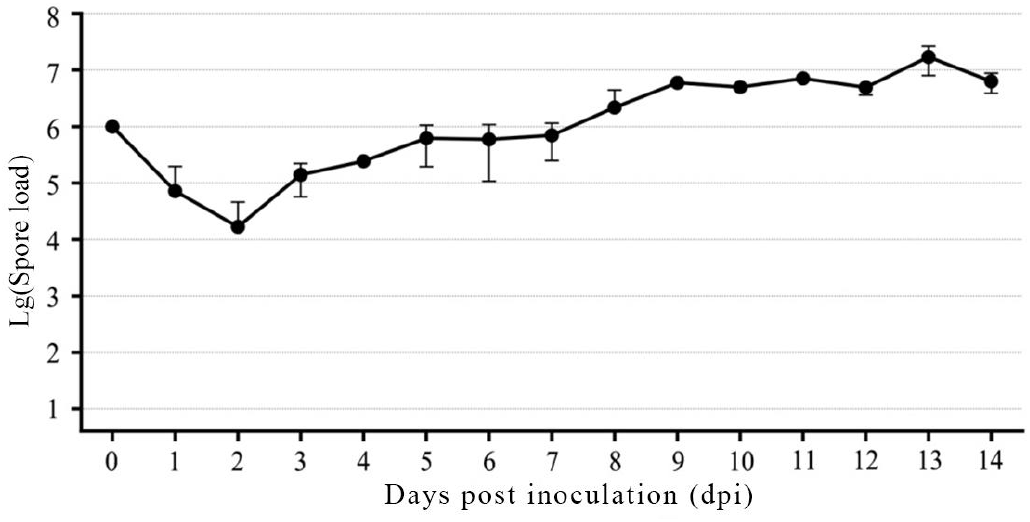
Spore load of *N. ceranae* in the *A. c. cerana* workers’ midguts after inoculation.

### 2.3 Sucrose solution consumption of *A. c. cerana* workers were elevated due to *N. ceranae* infection

The sucrose solution consumption of *A. c. cerana* workers was calculated, the result demonstrated that during the stage of 1 dpi–20 dpi, the sucrose solution consumption of workers in *N. ceranae*-inoculated groups was always higher than that of workers in un-inoculated groups, except 3 dpi, 17 dpi, and 20 dpi **(Figures 3)**; the average sucrose consumption of *N. ceranae*-inoculated and un-inoculated workers per day was 0.0357±0.0136 g and 0.0323±0.0066 g, respectively. In addition, the difference of sucrose solution consumption between these two groups at 4 dpi, 5 dpi, and 13 dpi was of significance **(Figure 3)**.

**Figure 3.**
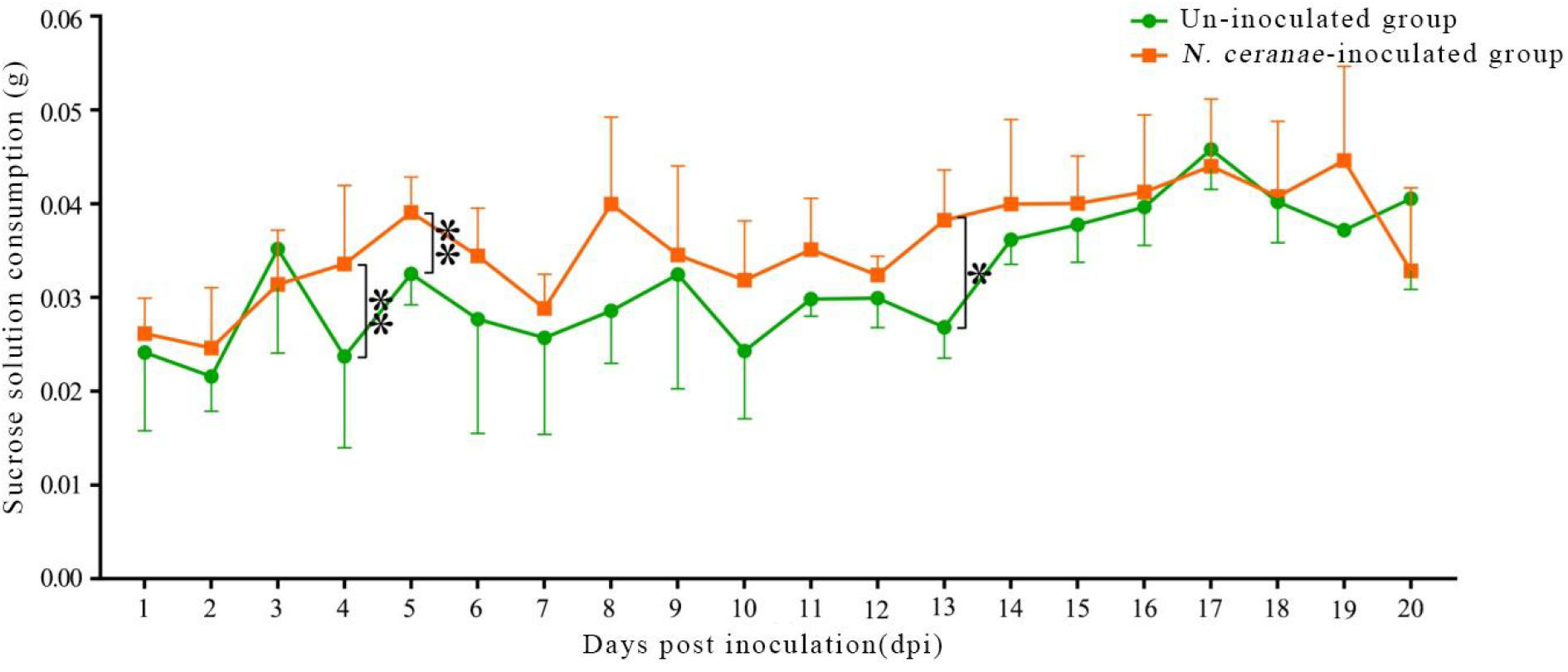
Sucrose solution consumption of *N. ceranae*-inoculated and un-inoculated workers.

### 2.4 *A. c. cerana* worker’s midgut epithelial cell structure was damaged by *N. ceranae* invasion

Microscopic observation of paraffin sections showed that darkly stained parasites were clearly detected in the midgut epithelial cells of *N. ceranae*-inoculated workers at 7 dpi-10 dpi, whereas no parasite was observed in those of un-inoculated workers **(Figure 4)**. Additionally, the fungal spores in the host cells gradually increased with the extension of infection time **(Figure 4)**. Moreover, the boundaries of un-inoculated host epithelial cells were intact and the darkly stained nucleus were clear, while the boundaries of midgut epithelial cells of *N. ceranae*-inoculated workers were blurred, the nucleus were almost disappeared, and the nucleic acid substances were diffused **(Figure 4)**.

**Figure 4.**
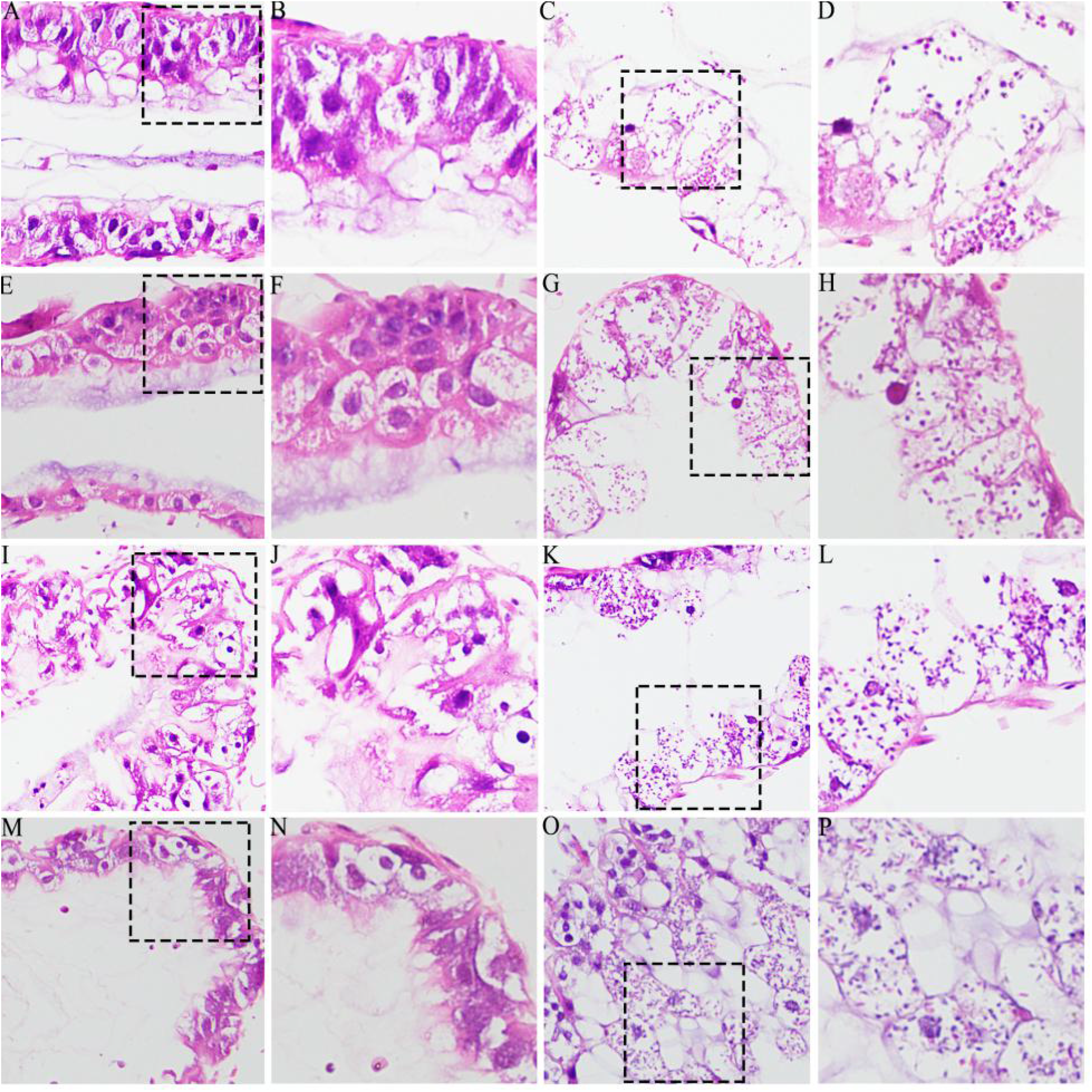
Microscopic observation of paraffin sections of un-infected and *N. ceranae*-infected *A. c. cerana* workers’ midguts. A-B: Midgut tissue of un-inoculated worker at 7 dpi without fungal spores; C-D: Midgut tissue of inoculated worker at 7 dpi with *N. ceranae* spores; E-F: Midgut tissue of un-inoculated worker at 8 dpi without fungal spores; G-H: Midgut tissue of inoculated worker at 8 dpi with *N. ceranae* spores; I-J: Midgut tissue of un-inoculated worker at 9 dpi without fungal spores; K-L: Midgut tissue of inoculated worker at 9 dpi with *N. ceranae* spores; M-N: Midgut tissue of un-inoculated worker at 10 dpi without fungal spores; O-P: Midgut tissue of inoculated worker at 10 dpi with *N. ceranae* spores. A, C, E, G, I, K, M, and O were microscopic fields under 200 times amplification, while B, D, F, H, J, N, and P were microscopic fields under 400 times amplification. Black dashed boxes show the selected region for observation under 200 times amplification.

### 2.5 *A. c. cerana* workers’ lifespan was influenced by *N. ceranae* challenge

Survival rate statistics indicated that the survival rates of workers in both *N. ceranae*-inoculated groups and un-inoculated groups at 1 dpi-5 dpi were pretty high, and there was little difference between the both groups **(Figure 5)**; the survival rates of both groups started to decrease at 5 dpi **(Figure 5)**; during the stage of 5 dpi-20 dpi, the survival rate of workers in *N. ceranae*-inoculated groups was always lower than that in un-inoculated groups, and the difference between these two groups was significant at 11 dpi-20 dpi**(Figure 5)**.

**Figure 5.**
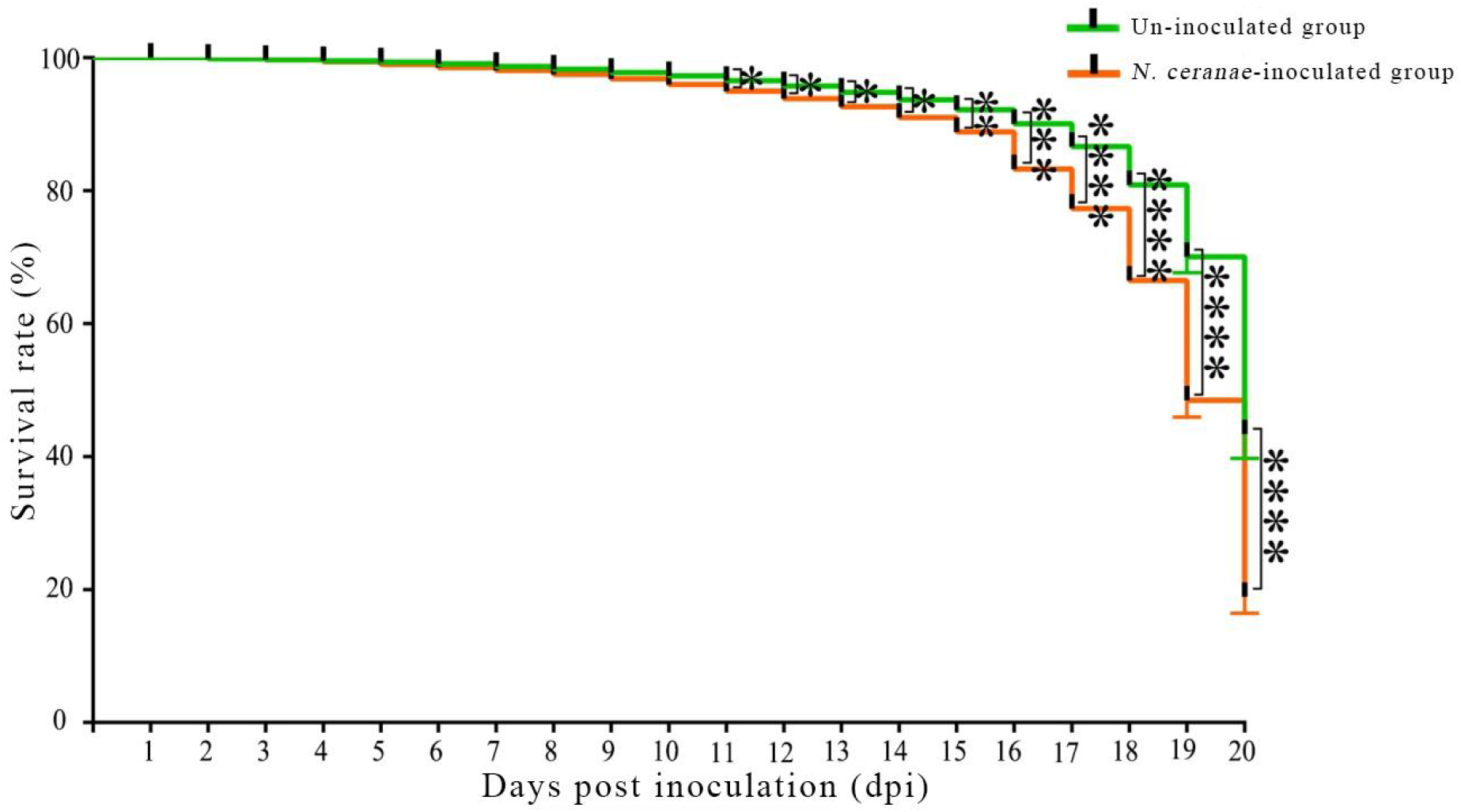
Survival rate of *A. c. cerana* workers after un-inoculation and inoculation with *N. ceranae* spores.

## 3. Discussion

Although *N. ceranae* originates from eastern honeybee, current understanding of their interaction is pretty limited, especially the *N. ceranae* infection. Previously work suggested both western honeybee and eastern honeybee workers could be effectively infected by artificial inoculation with *N. ceranae* spores [10-11]. In the present study, we investigated the impact of *N. ceranae* challenge on the sucrose solution consumption, midgut epithelial cell structure, and lifespan of *A. c. cerana* workers, the results demonstrated that the host sucrose solution consumption was elevated, the host epithelial cell structure was damaged, and the host lifespan was decreased, providing a solid experimental basis for further study on *N. ceranae* infection and microspoidian-eastern honeybee interaction.

Several previous studies were conducted to survey the spore load during the *N. ceranae* infection of *Apis mellifera* workers. Based on microscopy and molecular approaches, Huang et al. discovered that the spore load linearly elevated from 12 dpi to 20 dpi with *N. ceranae* spores [12]. Charbonneau and colleagues detected that the spore load of *N. ceranae* continuously increased in A. mellifera workers from 7 dpi to 14 dpi with fungal spores [13]. Li’s group measured the infection levels of *N. ceranae* in emergent bees, nurses, and foragers of *A. mellifera*, and found that the spore load and infection prevalence of *N. ceranae* varied among these adult workers and increased as workers aged [14]. Chen et al. observed that the spore load of *N. ceranae* in *Apis mellifera* workers was highly negatively correlated with temperature but not humidity [15]. In this current work, we detected that the spore load of *N. ceranae* showed a continuous elevation during the stage of 2 dpi-13 dpi, suggestive of the continuous proliferation of *N. ceranae* in host cells after inoculation. Additionally, microscopic observation of paraffin sections of *N. ceranae*-inoculated workers’ midguts demonstrated that the number of microsporidia in the epithelial cells increased during 7 dpi-10 dpi, further indicating the continuous multiplication of *N. ceranae* invading *A. c. cerana* workers. In detail, the spore load of *N. ceranae* in the *A. c. cerana* workers’ midguts decreased significantly during 1 dpi-2 dpi, and then continued to increase among 2 dpi-5 dpi, thereafter it slightly decreased during 5 dpi-6 dpi and continuously elevated among 6 dpi-9 dpi. This indicated that after entering the host midgut, the *N. ceranae* spores initiated the polar filament ejection mechanism, the polar tube punctured the epithelial cells and the inside infective sporoplasm was transferred into the host cells; the substances and energy in host cells were employed by *N. ceranae* to promote fungal proliferation, forming more and more progeny spores to complete the first round of proliferation, which were finally released to infect adjacent normal cells. It’s speculated that a life cycle of *N. ceranae* in the midgut epithelial cells of *A. c. ceranae* workers is approximately four days, similar to that of *N. ceranae* infecting *A. mellifera* worker’s midgut epithelial cells. This finding offers an experimental basis for further investigation of *N. ceranae* infecting eastern honeybee workers, such as functional study on fungal virulence factor-associated genes and development of RNAi-based control strategy. Here, on the basis of microscopic observation of paraffin sections, darkly stained microsporidia were clearly detected in the midgut epithelial cells of *N. ceranae*-inoculated workers during 7 dpi-10 dpi, and the boundaries of midgut epithelial cells of *N. ceranae*-inoculated workers were blurred and the nucleus were almost disappeared, with the nucleic acid substances dispersed within host cells; whereas the boundaries of un-inoculated workers’ midgut epithelial cells were intact. This suggests that the *N. ceranae* infection damages the structure of midgut cells of *A. c. cerana* workers.

Compared with un-infected *A. mellifera* workers, the sucrose solution consumption of *N. ceranae*-infected workers was significantly increased, reflecting the strong energy stress caused by *N. ceranae* invasion [16]. In this current work, it’s noted that the average sucrose solution consumption of *N. ceranae*-inoculated workers was always higher than that of un-inoculated workers during the stage of 1 dpi–20 dpi, except for 3 dpi, 17 dpi, and 20 dpi. The result indicates that the microsporidian infection also results in energy stress for *A. c. cerana* workers, which may lead to profound influence on host physiology such as cell growth and longevity.

Previous documentations showed that microsporidian infection significantly increased the mortality of both *A. mellifera* and *A. cerana* workers [7,17]. Paris et al. [16] assessed the survival rate and eating behavior of western bee workers infected by *N. ceranae* for 1–22 days, and found that the survival rate of infected workers was significantly lower than that of un-infected workers. In our previous study, we discovered that the mortality rate of *A. mellifera*-workers inoculated with *N. ceranae* spores gradually increased as time prolonged, and the mortality rate of *N. ceranae*-inoculated workers was significantly higher than that of un-inoculated workers at 7 dpi and 10 dpi [18]. Although *N. ceranae* was proved to be more infective for both *A. mellifera* and *A. cerana* workers than *N. apis* by assessing spore viability and infectivity [19], there was no convincing evidence regarding significantly different host mortality. Myrsini E et al. determined in their work that *N. ceranae* caused slightly higher host mortality compared to *N. apis*, but difference of virulence was non-significant [20]. Here, the survival rates of workers in both *N. ceranae*-inoculated groups and un-inoculated groups among 1 dpi-5 dpi were pretty high, implying that little influence on host longevity was caused by *N. ceranae* at the early stage of infection. In addition, during the stage of 5 dpi-20 dpi, the survival rate of *N. ceranae*-inoculated workers was always lower than that of un-inoculated workers, and the differences between them were significant during 11 dpi-20 dpi. The results demonstrate that the *N. ceranae* challenge exerts negative influence on the lifespan of *A. c. cerana* workers. In summary, it’s concluded that *N. ceranae* invasion can shorten the lifespan of both western honeybee and eastern honeybee workers.

## 4. Materials and Methods

### 4.1. Bee and microsporidian

*Nosema*-free workers of *A. c. cerana* were gained from colonies in the teaching apiary of College of Animal Sciences (College of Bee Science), Fujian Agricultural and Forestry University. No Varroa was observed during the whole experiment, and there was no specific band amplified from *N. apis, N. ceranae*, and several viruses such as SBV, DWV, IAPV, BQCV, KBV, and CBPV on the basis of RT-PCR using corresponding specific primers [9].

*N. ceranae*-infected workers of *A. c. cerana* were obtained from an apiary located in Minhou County, Fuzhou City, China. Clean spores of *N. ceranae* were previously prepared using a discontinuous density gradient Percoll method [9].

### 4.2 Experimental inoculation and artificial rearing of workers

Before artificial inoculation, the prepared fungal spores were subjected to microscopic detection with an optical microscope and PCR examination using a pair of specific primers for *N. ceranae* (F: CGGATAAAAGAGTCCGTTACC; R: TGAGCAGGGTTCTAGGGAT) previously described by Chen et al. [21].

Inoculation and rearing of *A. c. cerana* workers were performed following our previously established protocol [1,8]. Briefly, (1) workers (n=35) in treatment group at 24 h after emergence were each inoculated with 5 μL of 50% (w/v) sucrose solution containing *N. ceranae* spores (2×10^8^ per mL), while those (n=35) in control group were each inoculated with 5 μL of sucrose solution without microsporidian spores; (2) the *N. ceranae*-inoculated workers in plastic cages were reared in an incubator under the condition of 34±0.5 °Cand 60% ∼ 70% RH, whereas un-inoculated workers were reared in another incubator under the same conditions; (3) The honeybees were fed *ad libitum* with a feeder containing 4 mL of 50% (w/v) sucrose solution, and the feeders were replaced daily across the whole experiment; everyday, each cage was carefully checked and the dead worker bees were removed. There were three biological replicas of this experiment.

### 4.3 Measurement of spore load in workers’ midguts

According to the method described by our team [22], the spore load in workers’ midguts during the *N. ceranae* infection process was continuously counted. In brief: (1) after inoculation with *N. ceranae*, a single worker in treatment group was taken from the cage every day, and the midgut was then selected using a clean ophthalmic forceps and transferred into a sterile EP tube; (2) next, 200 μL of sterile water was added into the EP tube followed by full grinding with an automatic high-throughput tissue grinder (Meibi, Zhejiang, China); (3) 800 μL of sterile water was added to the grinding liquid and then fully mixed by repeated shaking; (4) ultimately, by using a micropipette, 100 μL of the aforementioned solution was carefully observed using a hemocytometer plate (Qiujing Shanghai, China) under a an optical microscope (CSOIF, Shanghai, China) followed by spore counting.

### 4.4 Detection of host sucrose solution consumption

Daily consumption of sucrose solution by each worker in *N. ceranae*-inoculated group or un-inoculated group was analyzed according to the described method as before [23-24]. The feeder containing 4 mL of sucrose solution was weighed, and the weight was recorded as A. To avoid the error due to repeated change of feeders, the feeder was weighed every 24 h thereafter, and the weight was recorded as B. The sucrose solution with the feeder was replaced daily. The sucrose solution consumption per worker at each day was calculated following the formula: (B-A)/number of survivors in the *N. ceranae*-inoculated group (or un-inoculated group).

### 4.5 Paraffin section, HE staining, and microscopic observation of workers’ midgut tissues

At 7 dpi-10 dpi, workers’ midguts in *N. ceranae*-inoculated group and un-inoculated group were respectively dissected out and fixed with 4% paraformaldehyde. By using an embedding center (Junjie, Wuhan, China) and a microtome (Leica, Nussloch, Germany), paraffin sections of midguts were stained with hematoxylin eosin (HE) stain by Shanghai Sangon Biological Engineering Co. Ltd., and then detected under an optical microscope with digital camera (SOPTOP, Shanghai, China).

### 4.6 Survival rate of workers

Following the protocol mentioned above, *A. c. cerana* workers were inoculated with *N. ceranae* spores and artificially reared; in another group, workers were fed with sucrose solution free of fungal spores. After inoculation, dead workers in both *N. ceranae*-inoculated group and un-inoculated group were recorded daily until 20 days post inoculation (dpi), followed by survival rate calculation and data analysis using GraphPad Prism software (GraphPad Company, USA).

### 4.7 Statistics

Statistical analysis and plotting of the data were performed using SPSS software (IBM Company, USA) and GraphPad Prism 7.00 software. Data were presented using mean ± standard deviation (SD), and the significance of sucrose solution consumption in the treated control group was analyzed using the two-tailed test in the Student’s *t* test. Log-rank (Mantel-Cox) test was used to analyze the survival rate of *A. c. cerana* workers. *P*<0.05 was considered statistically significant difference, and *P*<0.01 was considered highly significant difference.

## 5. Conclusions

In this current work, we performed in-depth investigation of the microsporidian infection and the impact of *N. ceranae* invasion on the host physiology, the results together indicate that the quantity of fungal spores continuously elevated with the microsporidian multiplication, leading to energetic stress for workers and host cell structure damage, which further negatively affected the host lifespan. Our findings provide a solid foundation not only for exploring the molecular mechanism underlying *N. ceranae* infection but also for investigating the interaction between *N. ceranae* and eastern honeybee workers.

## Author Contributions

R.G. and D.C. designed this research; Q.L., M.S. and X.F. contributed to the writing of the article; Q.L., M.S., X.F., W.Z., D.Z., Y.H., Z.W., K.Z., K.Y., H.Z., Y.S. and Z.F., conducted experiments and data analyses. R.G. and D.C. supervised the study and preparation of the manuscript.

## Acknowledgments

We thank all editors and reviewers for their helpful and constructive comments.

